# Transcriptome pyrosequencing of abnormal phenotypes in *Trypanosoma cruzi* epimastigotes after ectopic expression of a small zinc finger protein

**DOI:** 10.1101/002170

**Authors:** Gaston Westergaard, Marc Laverrière, Santiago Revale, Marina Reinert, Javier G. De Gaudenzi, Adriana Jäger, Martin P. Vazquez

**Affiliations:** Departamento de Fisiología y Biología Molecular y Celular, Facultad de Ciencias Exactas y Naturales, University of Buenos Aires, Argentina; Instituto de Investigaciones Biotecnológicas (IIB-INTECH), Universidad Nacional de San Martin, Buenos Aires, Argentina; Instituto de Agrobiotecnología Rosario (INDEAR), Rosario, Santa Fe, Argentina

## Abstract

The TcZFPs are a family of small zinc finger proteins harboring WW domains or Proline rich motifs. In *Trypanosoma brucei,* ZFPs are involved during stage specific differentiation. TcZFPs interact with each other using the WW domain (ZFP2 and ZFP3) and the proline rich motif (ZFP1). The tcZFP1b member is exclusive to *Trypanosoma cruzi* and it is only expressed in trypomastigote stage. We used a tetracycline inducible vector to express ectopically tcZFP1b in the epimastigote stage. Upon induction of tcZFP1b, the parasites stopped dividing completely after five days. Visual inspection showed abnormal distorted-morphology (monster) cells with multiple flagella and increased DNA contents. We were interested in investigate global transcription changes occurred during the generation of this abnormal phenotype. Thus, we performed RNA-seq transcriptome profiling with a 454 pyrosequencer to analyze the global changes after ectopic expression of tcZFP1b. The total mRNAs sequenced from induced and non-induced control epimastigotes showed, after filtering the data, a set of 70 genes having equal or more than 3X fold change upregulation, while 35 genes showed equal or more than 3X fold downregulation. Interestingly, several trans-sialidase-like genes and pseudogenes were upregulated along with several genes in the categories of amino acid catabolism and carbohydrate metabolism. On the other hand, hypothetical proteins, fatty acid biosynthesis and mitochondrial functions dominated the group of downregulated genes. Our data showed that several mRNAs sharing related functions and pathways changed their levels in a concerted pattern resembling post-transcriptional regulons. We also found two different motifs in the 3′UTRs of the majority of mRNAs, one for upregulated and other for downregulated genes

## Introduction

Trypanosomes are intriguing organisms in many aspects of their biology. In fact, they emerge as the paradigm to “the exception of the rule” in the eukaryotic lineage during the last decade. However, what was once considered “rare and exceptional” in these organisms was later shown to be more common than previously thought in the eukaryotic kingdom such as the mRNA trans- splicing or RNA editing processes [1].

Trypanosomes life cycles alternate between a mammalian host and an invertebrate vector. The adaptation to these two disparate environments requires a fine-tuning temporal control of gene expression and significant changes in the expression patterns of several genes. Notably, this regulation occurs almost entirely at the post-transcriptional level and typical polymerase II promoters are absent [1,2].

In accordance, genome organization and expression in trypanosomatids is unusual. Transcription of protein coding genes is not regulated at the level of transcription initiation by RNA polymerase II and their genes are organized into densely packed units with relatively short intergenic regions and mostly devoid of introns. In this way, gene expression is polycistronic and controlled mainly by post-transcriptional processes [1,3,4].

Polycistronic units contain unrelated genes that are co-transcriptionally processed to individual mRNAs by two coupled reactions controlled by the intergenic regions: 5′trans-splicing and 3′polyadenylation. Thus, the cis-acting sequences and trans-acting factors controlling the post-transcriptional gene expression in these parasites are extremely important [5].

Several trans-acting factors involved in these processes were identified, such as RRM (RNA Recognition Motif) containing proteins, PUF proteins and CCCH containing proteins, comprising a group generally known as RBPs (RNA Binding Proteins) [6,7,8].

RBPs have shown to regulate mRNA abundance in trypanosomes through a number of cis-acting sequences, most notably AU rich elements (AREs) for RRMs and UGUR core elements for PUF proteins, both type of sequences present in the 3′end of their target transcripts [9,10].

Genome-wide analysis of these proteins in the Tritryps (*Leishmania major*, *Trypanosoma brucei* and *Trypanosoma cruzi*) showed that they contain between 75 (*T. brucei*) and 139 (*T. cruzi*) RRM-type proteins [6].

They also contain 10 different PUF proteins and between 48 and 54 CCCH-type proteins encoded in their genomes [10].

The majority of the CCCH-type proteins are unique to the Tritryps and they share a core of 39 proteins in common with differences due to either loss or gain of a single gene-by-gene duplication.

An important set of CCCH-type proteins was previously studied in *T. brucei* and *T. cruzi*: the tbZFPs and tcZFPs [11,12].

The tbZFPs are a group of two small proteins, tbZFP1 (101 residues) and tbZFP2 (139 residues). The tbZFP2 also contains a WW domain characteristic of protein-protein interactions with proline-rich motifs. Genetic perturbation assays provided the evidence that the two tbZFPs can regulate differentiation and morphogenesis in *T. brucei*. Overexpression of tbZFP2 generated a posterior extension of the microtubule corset, a mechanism responsible for kinetoplast repositioning during differentiation [13] and RNAi mediated knockdown of tbZFP2 severely compromised differentiation from bloodstream to procyclic forms. It was also shown that tbZFP1 is enriched through differentiation to procyclic forms.

Later on, a new tbZFP was described, tbZFP3 (CCCH and WW domains), which enhances development among life cycle stages in T. brucei. Ectopic expression of tbZFP3 in the insect stage of the parasite produced elongated forms (nozzle phenotype) typical of induced differentiation; while ectopic expression in the bloodstream stage enhanced differentiation by upregulating EP procyclin protein expression [14].

In *T. cruzi*, four tcZFPs were described, tcZFP1a, tcZFP1b, tcZFP2a and tcZFP2b. The tcZFP1 proteins present CCCH and proline-rich motifs, whereas tcZFP2 proteins present the CCCH motif and a WW domain. Interestingly, tcZFP2s engage in protein-protein interactions with tcZFP1s via the WW domain and the proline-rich motif respectively. Another interesting thing to note is that tcZFP1b is *T. cruzi* specific due to a partial gene duplication event that conserved the central core region of the tcZFP1a protein and diverged in the N- and C-terminal parts of the protein [11].

Different tcZFPs are expressed in different life-cycle stages, thus allowing for a modularization of the protein-protein interactions and it is proposed that this may allow the control of expression of distinct cohorts of genes in different life-cycle stages in a form of post-transcriptional operonic regulation [15].

In this work, we focus our attention in tcZFP1b, the *T. cruzi* specific protein. Since we previously showed that tcZFP1b mRNA was absent in epimastigotes (the insect stage) and was only detectable in trypomastigotes (the bloodstream stage), we devised that through the analysis of its ectopic expression in epimastigotes we could gain insight into the function of this protein.

Overexpression of tcZFP1b translates into a cell cycle arrest. An RNA-seq transcriptome profiling comparison of non-induced and induced cells showed several mRNAs of related functions changing in a concerted post-transcriptional pattern resembling post-transcriptional regulons.

## Results

### Ectopic overexpression of tcZFP1b in epimastigotes (insect stage)

Initial attempts to overexpress tcZFP1b in epimastigotes using constitutive overexpressing vectors such as pTREX [16] were unsuccessful for the selection of stable transgenic populations. In fact, parasites stop dividing two weeks after transfection and died systematically after various attempts. Visual inspection showed several monster cells (aberrant phenotype with multiple flagella and possible multiple nuclei or kinetoplasts) few days before parasites died. On the other hand, transfection with unrelated genes such as eGFP or other *T. cruzi* genes were successful [16].

Thus, we decided to clone tcZFP1b and the control gene eGFP in the inducible vector pTcINDEX [17] (Fig. 1A). After transfection of culture epimastigotes and clonal selection of strains, parasites were growing normally. After addition of tetracycline to induce gene expression, eGFP transgenic parasites grew normally for six days before reaching a plateau. In contrast, tcZFP1b transgenic parasites showed a marked decrease in cell counts by day three and eventually stop dividing by day four (Fig. 1A). Control tcZFP1b parasites without tetracycline addition grew normally until reaching a plateau by day six (Fig. 1A).

**Figure 1.**
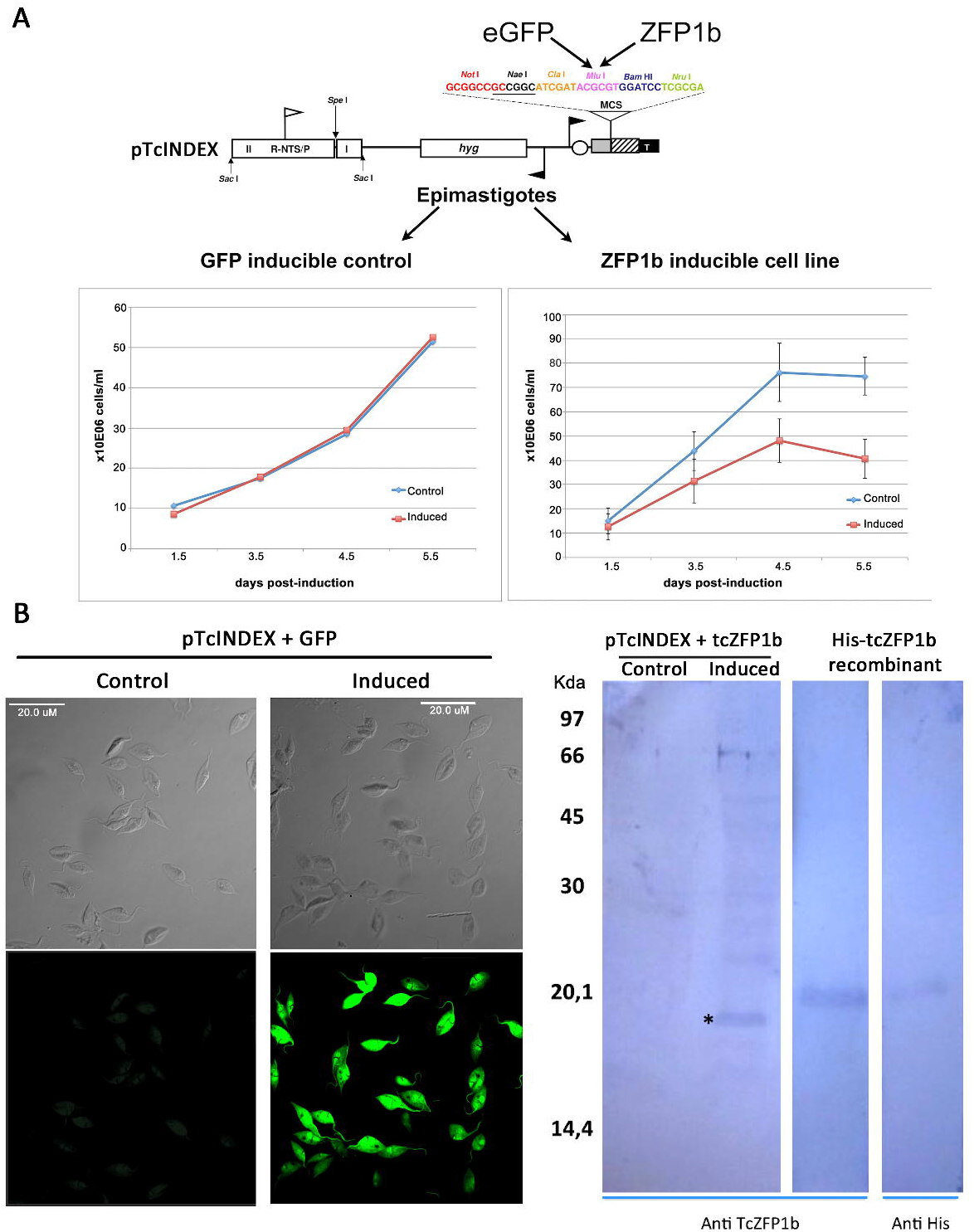
Inducible overexpression of tcZFP1 in epimastigotes. **A:** Upper panel, schematic representation of pTcINDEX, as appeared in [17], with the multiple cloning site used to introduce the eGFP and tcZFP1b genes. Both vectors were transfected into epimastigotes to generate independent cell lines. The eGFP was used a as control to test overexpression and leakage. Lower panel, growth curves for GFP and tcZFP1b cell lines induced with tetracycline and non-induced (control). **B:** Left panel, overexpression test of GFP using confocal microscopy. Results showed no leakage for control (non-induced) and strong induction after tetracycline addition (induced). Right panel, Western blot confirmation of tcZFP1b overexpression using a tcZFP1b specific mouse antiserum. Antibodies specificity was tested against a His-tcZFP1b recombinant protein produced in bacteria.

Visual analysis of induced expression and background leakage from the system was done by confocal microscopy for eGFP and by western blot for tcZFP1b. Results showed high levels of expression upon induction, while the system showed no background leakage as measured by day four (Fig. 1B)

### Monster cells in tcZFP1b transgenic cell line are arrested in cytokinesis and G2 phase of the cell cycle

Presence of abnormal shaped (Monster) cells was observed in the culture of tetracycline induced tcZFP1b transgenic epimastigotes.

Confocal microscopy inspection showed that epimastigotes were arrested at an early step of cytokinesis. Parasites presented two flagella but no cytoplasmic division. In contrast, non-induced control parasites were often seen with two flagella in opposite orientation and proceeding with final steps of cell division (anti-tubulin staining, Fig. 2A). By using propidium iodide staining for DNA, it was clearly seen that monster cells also contained two nuclei and two kinetoplasts (2N2 K content) (PI staining, Fig. 2B).

**Figure 2.**
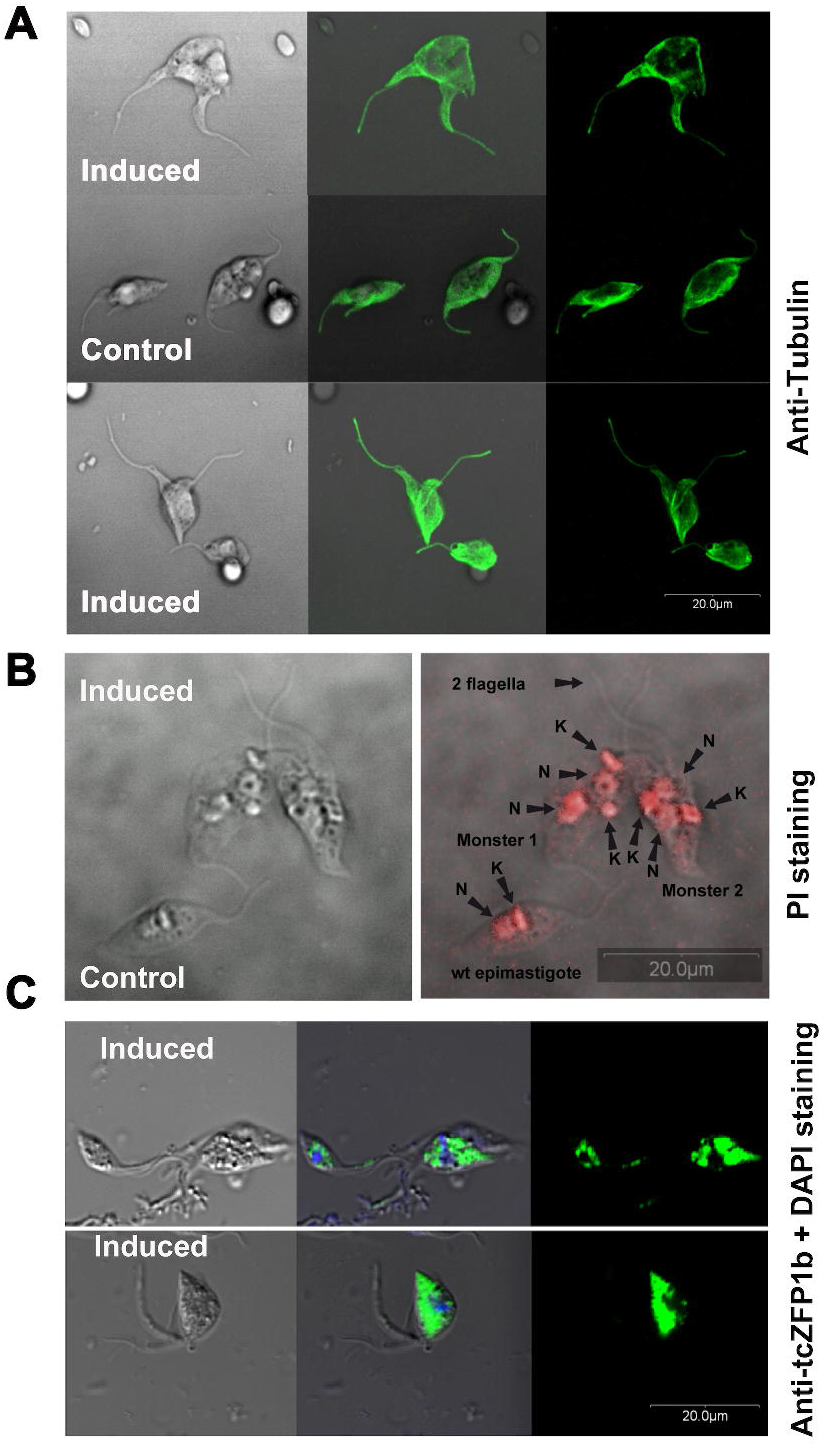
Monster cells appeared upon overexpression of tcZFP1b in epimastigotes. **A:** Confocal microscopy images of tcZFP1b induced and control (non-induced) epimastigotes detected using anti-tubulin specific antibodies (FITC conjugated anti-mouse antibody). Induced cells showed monster phenotypes with arrested cytokinesis while non-induced cells showed wild type phenotypes and parasites proceeding with typical cell division. **B:** Confocal microscopy images of epimastigotes stained with propidium iodide (PI) for detection of DNA. Induced and control parasites were mixed 50:50 in one slide for visualization and direct comparison of wild type and monster phenotypes. N, nucleus; K, kinetoplast DNA. **C:** Confocal microscopy images of induced epimastigotes detected using anti-tcZFP1b specific polyclonal serum (FITC conjugated anti-mouse antibody) and DAPI staining for DNA. The central panels show merged images of DIC (differential interface contrast), FITC detection and DAPI staining. Overexpressed tcZFP1b is distributed in the cytoplasm excluding nucleus and kinetoplast.

This was confirmed by FACS analysis (Fig. S1A). A significant number of cells that left G1 were arrested at G2/M tcZFP1b induced cells respect to non-induced control cells. Even a considerable number of epimastigotes presented 4N content compared to control. DAPI staining of DNA from the same samples confirmed multiple nuclei and kinetoplasts in cells with multiple flagella (Fig. S1B).

The monster cells phenotypes were almost identical to those obtained by treating epimastigotes with taxol (an anti-tumoral agent and stabilizer of microtubules) [18]. This fact was strongly indicative that overexpression of tcZFP1b led to a cytokinesis arrest similar to that of taxol treatment [18].

Analysis of the induced expression of tcZFP1b in the same samples was confirmed by confocal microscopy and indirect immunofluorescence using anti-tcZFP1b serum. Results showed that tcZFP1b was highly expressed upon induction and that the protein is distributed in a particulate form over the cytoplasm excluding nucleus and kinetoplast (Fig. 2C)

To gain insights into the induced cell cycle arrest, we performed Transmission Electron Microscopy (TEM) in the tcZFP1b induced samples. The micrographs showed, as expected, multiple nucleus and kinetoplast per cell and also showed evidence of multiple flagellum (Fig. 3). Interestingly, we detected cases with two kinetoplasts and four basal bodies (asterisks in Fig. 3B) suggesting basal body duplication without kinetoplast duplication, indicative of the start of another round of cell division without cytokinesis. It is also interesting to note the distribution of the chromatin within the nuclei. According to Elias et al. [19], the concentration of the chromatin in dense granules in the nuclear periphery attached to the envelope is indicative of cells being in the G2 phase of the cell cycle. A great majority of the cells showed this particular distribution of the chromatin (see Cr in Fig. 3).

**Figure 3.**
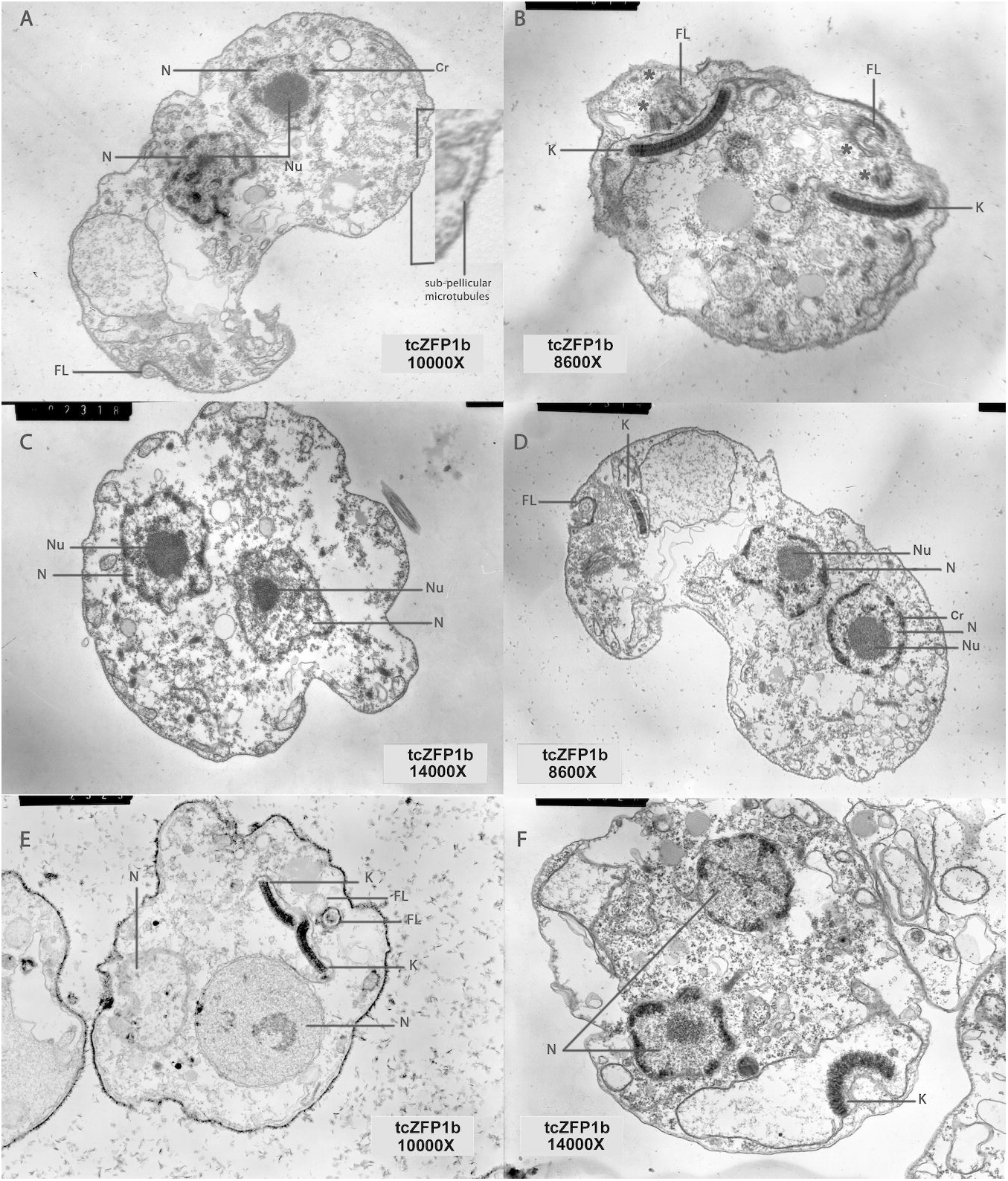
Monster epimastigotes arrested in G2-phase of the cell cycle upon overexpression of tcZFP1b. Six images of monster phenotypes obtained using Transmission Electron Microscopy (TEM). N, nucleus; Nu, nucleolus; K, kinetoplast; FL, flagellum; Cr, chromatin. Asterisks in image **B** denotes basal body duplications. Dense granules in the nucleus indicate chromatin attached to the envelope and it is indicative of the G2-phase of the cell cycle.

### Changes in the epimastigote transcriptome profile upon overexpression of tcZFP1b as determined by 454 pyrosequencing

Ectopic expression of tcZFP1b produced aberrant phenotypes and cytokinesis arrest. Since tcZFP1b is an RBP and trypanosomes control gene expression post-transcriptionally, we decided to investigate global changes in the expression profile using RNA-seq analysis.

Samples were taken in duplicate at 70 hours after tetracycline addition when cell replication arrest had begun (Induced) or no-addition (Control) (Fig. S2A).

The mRNA pyrosequencing produced a total of 233,310 reads for Control and 206,703 reads for Induced. The uniquely mapped reads used for the analysis of quantitative expression were 95,792 and 107,111 for each duplicate in control, while 73,490 and 112,570 were mapped for Induced (Fig. S2B). A total of 30,407 and 20,643 reads remained unmapped for Control and Induced, respectively.

Uniquely mapped reads were normalized by depth and gene length as indicated in Methods section. As a standard internal control, we measured the changes in gene expression of tcZFP1b after induction as well as the endogenous tcZFP1a. It was confirmed that tcZFP1b mRNA presented a 126X fold change upon induction while endogenous tcZFP1a expression was low and showed no changes (Fig. S2C). Importantly, it was also confirmed that leakage from pTcINDEX inducible vector was very low to non-existent.

To analyze coverage and bias in the 454 RNA-seq, we used three different genes as example: the internal control tcZFP1b (378 b), a downregulated gene (α-tubulin, 1318 b), and an upregulated gene (proline racemase, 1065 b) (Fig. S3). Coverage was very good in all cases as expected for the long read sequences of the 454 technology, even in low-medium expressed genes (i.e. proline racemase in control) (Fig. S3). The coverage presented a little bias to the 3′end, which was also expected for this methodology, although lower due to the mRNA fragmentation procedure instead of cDNA fragmentation (Fig. S3) [20].

To analyze global changes, we established a set of rules in order to look for the most prominent changes. First, we filtered the data set so that genes with at least five uniquely mapped reads in any condition were retained. This produced a data set of 2737 genes for analysis. Then, we filtered for those genes between 10 and 100 unique mapped reads that presented a 3X fold or more change. We also filtered for those genes with unique mapped reads between 100 and 1000 that presented a 2X fold or more change. In this way, we established a data set of 112 genes that match our criteria of regulation above threshold (Fig. 4). Under these conditions, a total of 73 genes were upregulated and 39 genes were downregulated.

**Figure 4.**
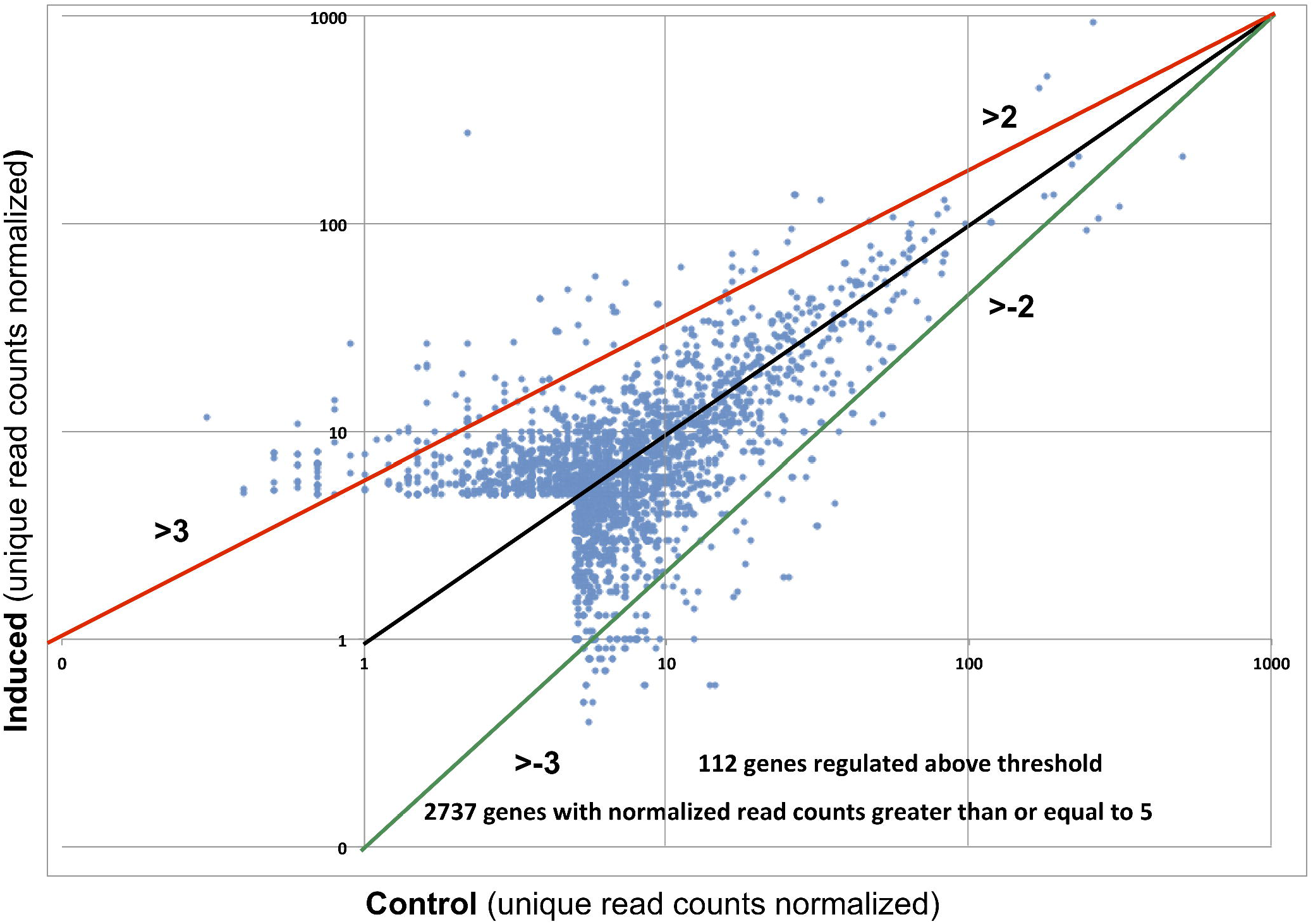
Transcriptome profile of induced versus control (non-induced) epimastigotes. Normalized unique read counts from induced and control sequenced RNA samples of epimastigotes were plotted against each other in a log scale. Each blue dot represents a different gene. Genes with equal or more than five normalized read counts were plotted (2737 genes). The red line represents the threshold of upregulated genes and the green line represents the threshold for the downregulated ones in the induced sample. Two different thresholds were considered as significant: a) 3 or more fold change in the range from 5 to 100 unique read counts; and b) 2 o more fold change in the range from 100 to 1000 unique read counts. A total of 112 genes changed expressions above these thresholds.

### Upregulated genes in induced versus control epimastigotes

Among the top 15 expressed genes in epimastigotes, the tyrosine aminotransferase (TAT), mucin TcSMUGS and hexose transporter genes were upregulated, while α-tubulin and prostaglandin F2α synthase genes were downregulated (Fig. 5). Interestingly, TAT genes, which were the top fourth and eighth expressed genes in the control epimastigote, became the two most expressed in the induced epimastigote (3.6X fold change) (Fig. 5).

**Figure 5.**
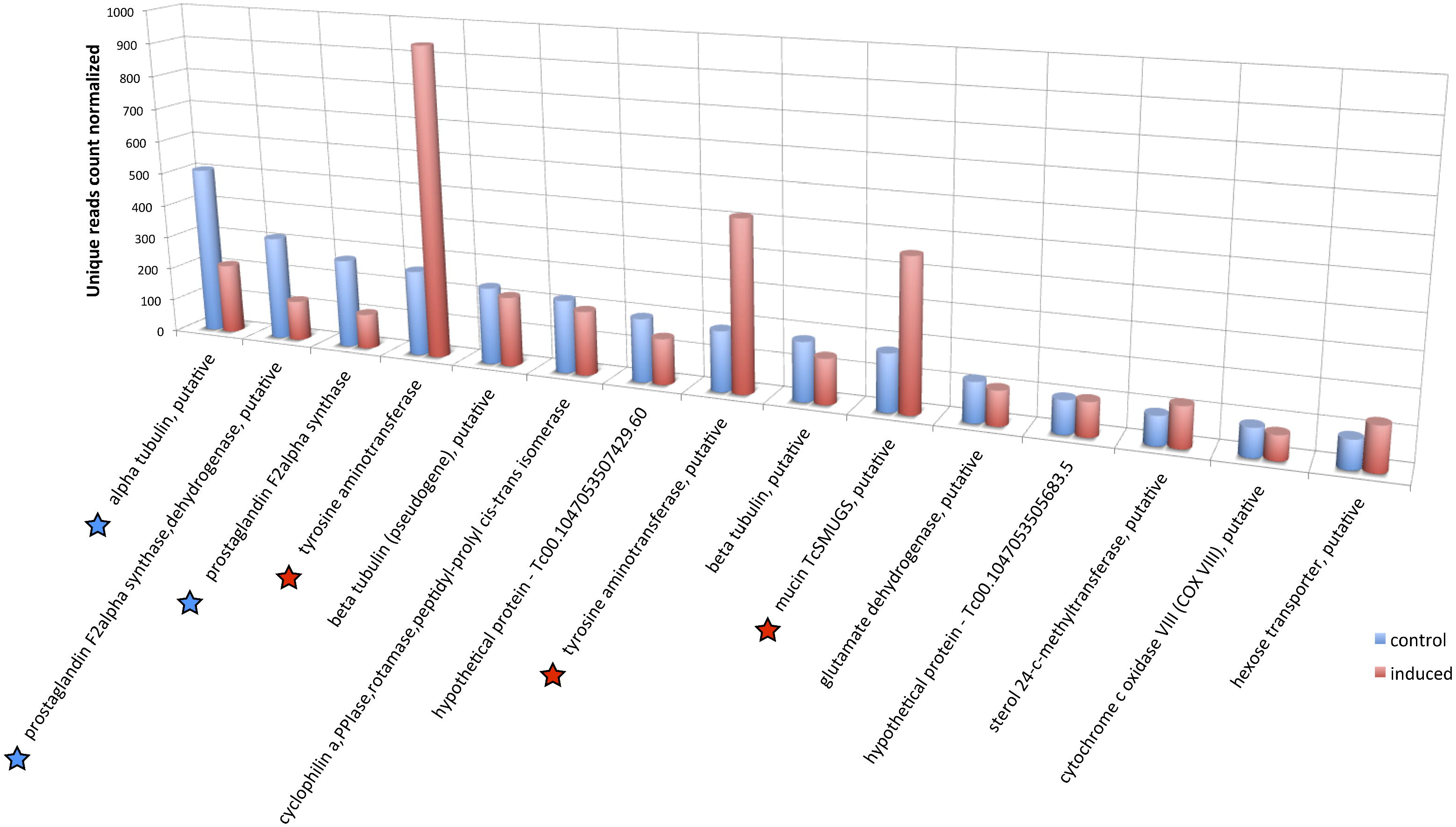
Expression profile of top 15 expressed genes in control versus induced epimastigotes. The top 15 expressed genes in the control (non-induced) RNA sample were compared with their expression in the induced RNA sample and the normalized unique read counts were plotted. A star below the gene description name indicates a significant change in mRNA levels between the two samples. A blue star denotes a downregulation and a red star an upregulation in the induced sample. Similar description names indicate similar genes from the two *T. cruzi* haplotypes reference genomes, esmeraldo and esmeraldo-like, of the CL-Brener strain.

Among the 3X fold change upregulated genes, we could establish five different categories: infectivity and differentiation, carbohydrate metabolism, amino acid metabolism, ribosomal function and hypothetical proteins (Fig. 6). Interestingly, the dominant top 20 upregulated genes belonged to the categories of infectivity and differentiation and amino acid metabolism.

**Figure 6.**
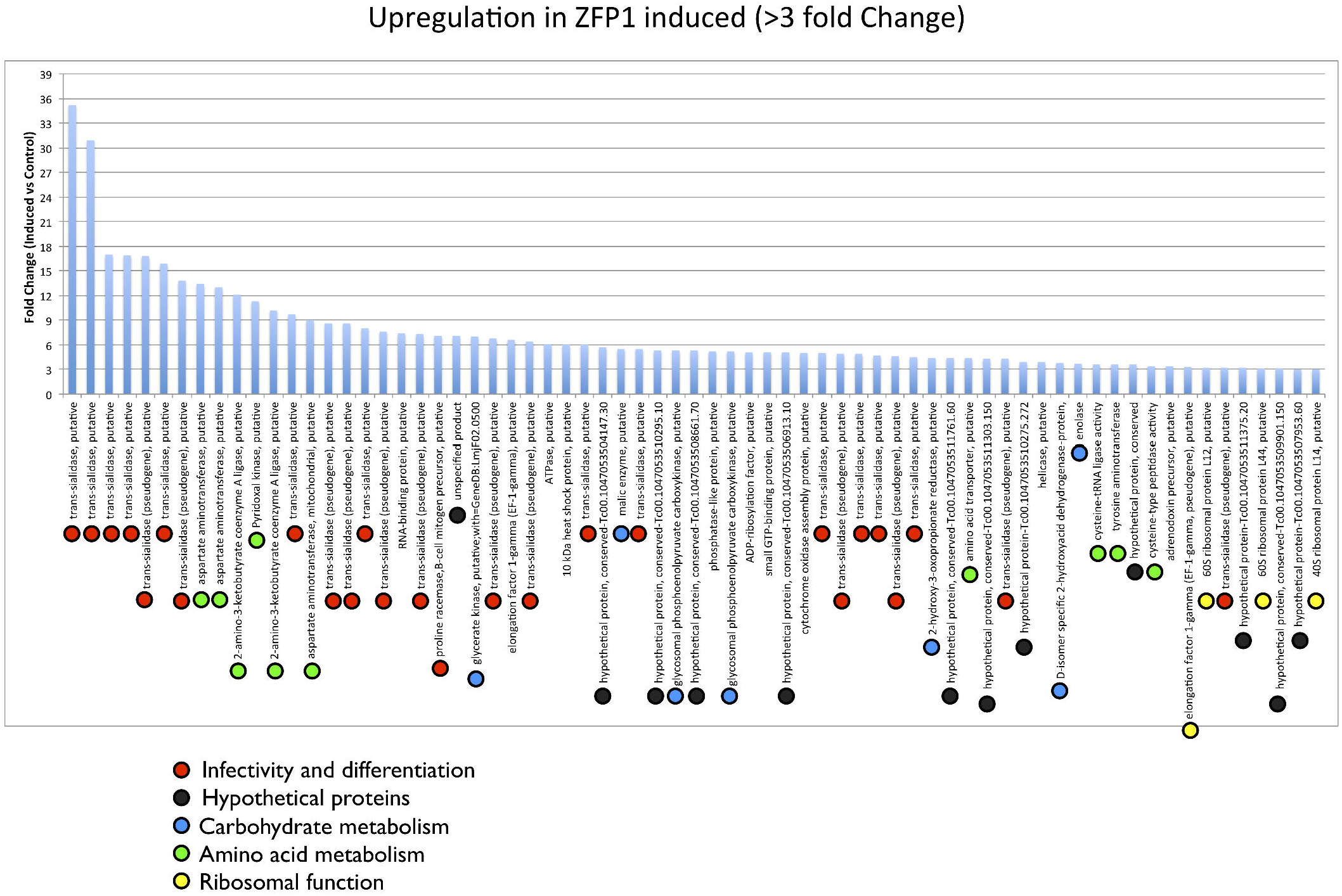
Upregulated genes in epimastigotes upon overexpression of tcZFP1b. The fold change (induced versus control) was plotted considering only those genes with changes equal or more than 3X. Upregulated genes were categorized using different colored dots as indicated. For proteins with predicted function, the description name is provided. For hypothetical proteins, the GeneDB number is provided

The infectivity and differentiation category is populated with a majority of trans-sialidase like family of genes, a family composed of hundreds of members, dispersed throughout all chromosomes and mostly expressed on the surface of trypomastigotes (bloodstream stage) [21]. Notably, several trans-sialidase like pseudogenes were also expressed and upregulated. Fold changes for trans-sialidase like members were from 35X to 3X (Fig. 6).

Another important upregulated gene was proline racemase (7.1X fold change), which was demonstrated to participate in differentiation from epimastigotes to trypomastigotes and to enhance infectivity of host cells (Fig. 6) [22,23].

The second important category was amino acid metabolism represented by the genes of aspartate aminotransferase (cytoplasmic and mitochondrial), 2-amino-3-ketobutyrate coenzyme A ligase and tyrosine aminotransferase. Their upregulation ranged from 13X to 3X fold change. The products of these genes participate in amino acid catabolism. Interestingly, they use pyridoxal phosphate as a cofactor and pyridoxal kinase was also one of the upregulated genes (11X fold change). Another interest correlation was the upregulation of an amino acid transporter gene (4.4X fold change).

The third important category was related to carbohydrate energy metabolism. The glycosomal phosphoenolpyruvate carboxykinase (5.3X fold change) and malate dehydrogenase (5.5X fold change) were known to contribute to ATP regeneration in the glycosome [24]. Moreover, D-2-hydroxy-acid dehydrogenase (3.8X fold change) was involved in the conversion of lactate to pyruvate and 2-hydroxy-3-oxopropionate reductase (4.4X fold change) was involved in the glyoxylate-dicarboxylate metabolism. The glycerate kinase gene was upregulated by 7X fold and was also related to glyoxylate-dicarboxylate metabolism (Fig. 6).

These processes were also linked to the glycine, serine, threonine metabolism described above to provide glyoxylate and hydroxipyruvate.

A minor group of genes related to ribosomal functions were upregulated such as elongation factor-1 gamma (6.6X fold), L14 (3.1X fold) and L44 (3X fold) ribosomal proteins. Additionally, 12 genes coding for hypothetical proteins were upregulated ranging from 7X to 3X fold change (Fig. 6)

It is important to note that all these upregulated genes are not located close to each other in the genome. Moreover, they are dispersed in different chromosomes in most cases.

### Downregulated genes in induced versus control epimastigotes

Among the top 15 expressed genes, it is worth to mention that the top three were downregulated: α-tubulin (2.4X fold) and the two genes for prostaglandin F2α synthase (2.6X and 2.5X fold) (Fig. 5). Interestingly, it was shown that prostaglandin F2α (PGF2) was mainly produced in fast dividing forms of the parasite (i.e epimastigotes) and it was lower in non-dividing forms or during stationary phase in culture [25]. Downregulation of PGF2 synthase correlated perfectly with the fact that induced epimastigotes stopped dividing at the time of sampling (Fig. S2).

Among the group of 3X fold downregulated genes, we established three different categories: hypothetical proteins, fatty acids biosynthesis and mitochondrial functions (Fig. 7). The hypothetical proteins dominated the group of downregulated genes in induced epimastigotes. This fact was drastically different from the situation with the upregulated genes. This precludes a comprehensive analysis of the downregulated functions. However, it is worth mentioning that almost 60% of the hypothetical proteins were conserved among the trypanosomes, while the other 40% were *T. cruzi* specific (Fig. 7).

**Figure 7.**
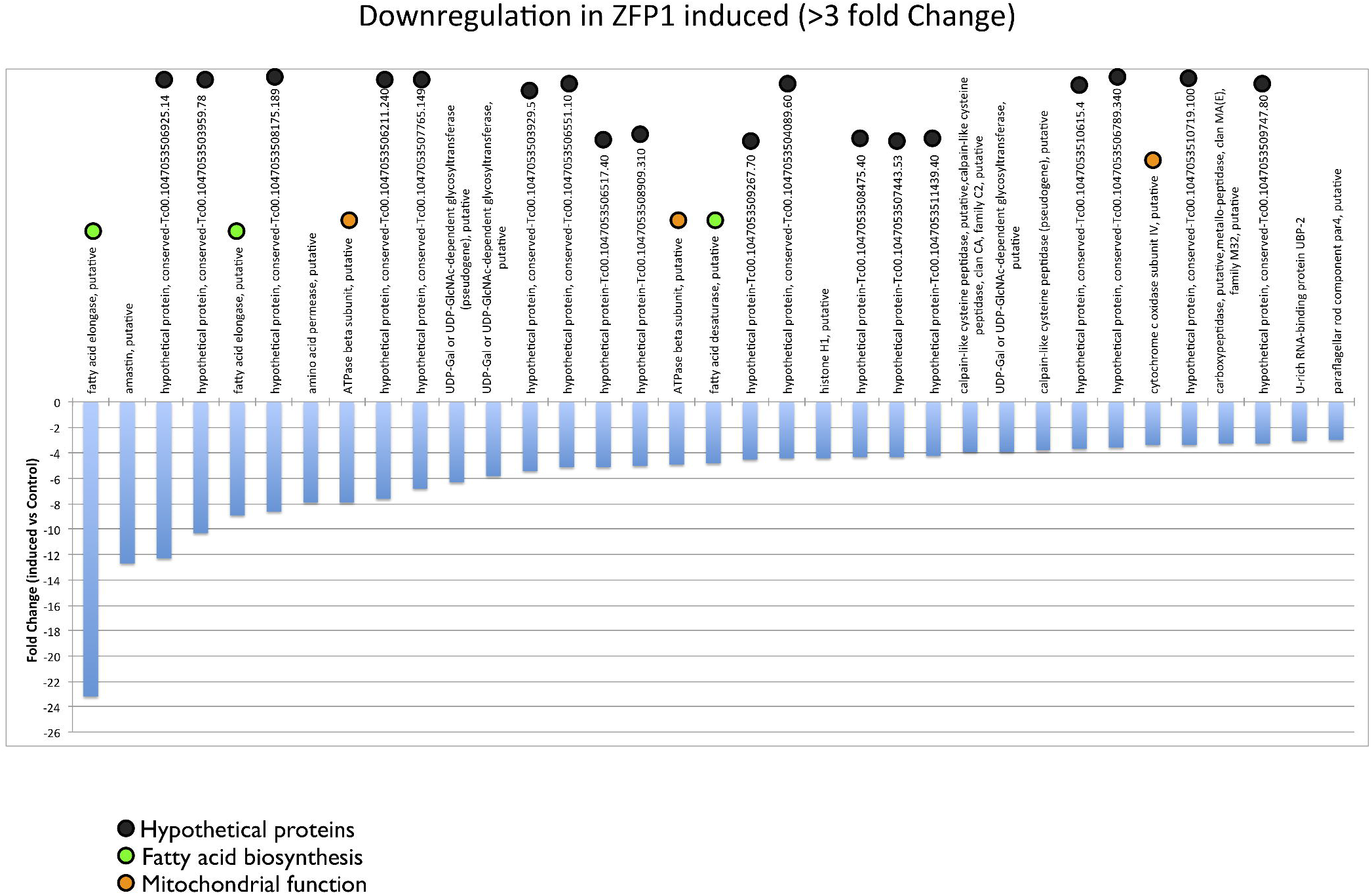
Downregulated genes in epimastigotes upon overexpression of tcZFP1b. The fold change (induced versus control) was plotted considering only those genes with changes equal or more than 3X. Downregulated genes were categorized using different colored dots as indicated. For proteins with predicted function, the description name is provided. For hypothetical proteins, the GeneDB number is provided.

Downregulated mRNAs with annotated functions involved an important group related to fatty acid biosynthesis, such as fatty acid elongase (9X fold) and fatty acid desaturase (5X fold). A possible downregulation of mitochondrial functions could be also suggested due to downregulation of ATPase beta subunit (8X fold) and cytochrome C oxidase subunit IV (3.5X fold).

Other downregulated genes include amastin (12.7X fold), which was known to be downregulated in trypomastigotes [26], an amino acid permease (7.9X fold), histone H1 (4.4X fold), the UBP-2 RNA binding protein (mRNA metabolism, 3.1X fold) and the paraflagellar rod component par4 (3X fold)

To validate the quantification analysis of RNA-seq, we performed qPCR of selected genes. We chose three upregulated genes (aspartate aminotransferase cytoplasmic and mitochondrial, and piridoxal kinase), four downregulated genes (α-tubulin, PGF2 synthase, ATPase beta subunit, and carboxipeptidase) and three genes with no expression changes (ribosomal protein L35a, gapdh, and Acyl carrier protein). The qPCR was performed in triplicate for each gene in control and induced samples. The results indicated a very good correlation with the RNA-seq data for all the genes tested (Fig. S4).

### Sequence motifs in upregulated and downregulated genes

To evaluate the presence of common sequence motifs in the 3′UTRs among the data set of 112 genes with changed expression above threshold, we selected a fixed window of 300 nt downstream the stop codon.

The bioinformatics analysis was done as described in the Methods section. We found two different high-confidence motifs in the data set: up-h12 and down-h12 (Fig. 8). The up-h12 motif presented a conserved core of UGuxxxGxGc (Fig. 8A), while down-h12 was a well-conserved AU-rich (ARE) element [6,9] with a conserved core UxUAU (Fig. 8B). Both motifs could be folded into stem-loop structures with the conserved cores exposed in the loops (Fig. 8).

**Figure 8.**
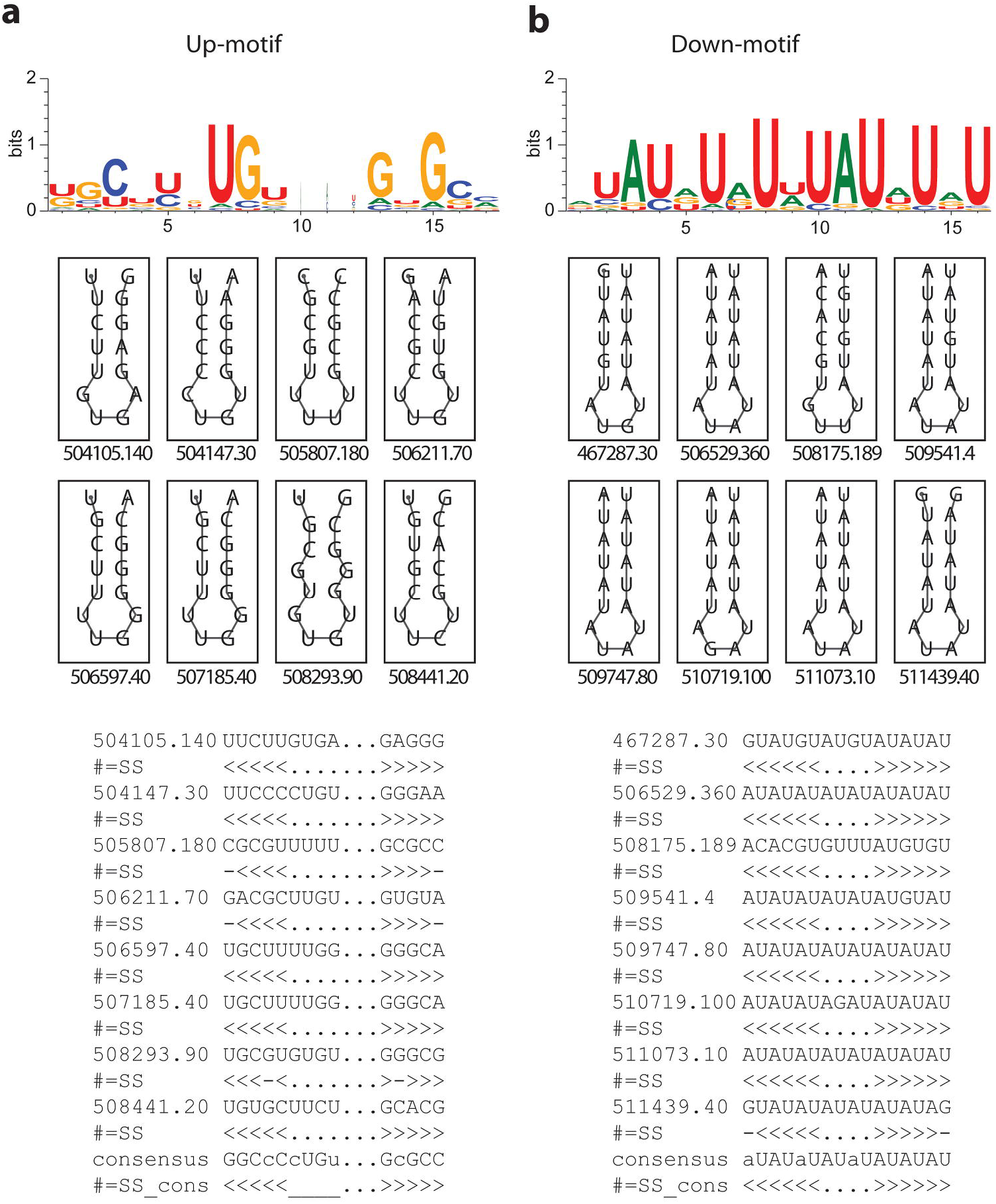
Common motifs in the 3′UTR of upregulated and downregulated genes in induced epimastigotes. **A:** Consensus motif found in upregulated genes. **B:** Consensus motif found in downregulated genes. Upper panel, sequence logo representations of consensus motifs. Middle panel, secondary structures for eight selected sequences representing the found motif, each one with their unique identifier of the GeneDB number below (http://tritrypdb.org). Lower panel, linear sequences for the selected 3’UTRs with their corresponding consensus. The selected 3’UTR for up-h12 are 504105.140, enolase; 504147.30, hypothetical protein, conserved; 505807.180, 2-hydroxy-3-oxopropionate reductase; 506211.70, RNA-binding protein; 506597.40, trans-sialidase; 507185.40, trans-sialidase (pseudogene); 08293.90, elongation factor 1-gamma (EF-1-gamma); 508441.20, glycosomal hosphoenolpyruvate carboxykinase. The selected 3’UTR for down-h12 are 467287.30, ATPase beta subunit; 506529.360, cytochrome c oxidase subunit IV; 508175.189, hypothetical protein, conserved; 509541.4, paraflagellar rod component par4, putative; 509747.80, hypothetical protein, conserved; 510719.100, hypothetical protein, conserved; 511073.10, fatty acid desaturase; 511439.40, hypothetical protein.

The motif up-h12 was found with coverage of 72% in the data set of upregulated genes and 38% in the downregulated genes. The motif down-h12 was found with coverage of 89% in downregulated and 27% in the upregulated genes (for the complete list of genes containing up-h12 and down-h12 and their positions in the 3’UTR see Table S2). Statistical significance was determined using a chi-square test by comparing the motifs against 115 groups composed by 50 randomly selected 3′UTR sequences. Results indicated that up-h12 could be found by chance in the data set with coverage of 45.49% (p < 0.01), while down-h12 could be found at random with coverage of 45.37% (p < 0.005). Thus, the presence of up-h12 and down-h12 were not random and has statistically significant correlations.

The identification of common sequence motifs in the 3′UTR of the up and down regulated mRNAs reinforces the idea of possible post-transcriptional regulons in *T. cruzi*.

### Discussion

Post-transcriptional regulation is highly dependent on RBPs to achieve a fine-tuning control of mRNA levels. This is particularly important in a parasite adapted to two disparate environments in different hosts encountering abrupt changes that occur in a timeframe of seconds.

Several RBPs were described in trypanosomes and they were shown to be involved in post-transcriptional regulation [6]. Accordingly, the RRM, PUF and zinc finger domains and motifs are expanded in these parasite genomes [7].

Within the ZFP family, tcZFP1b is exclusive to *T. cruzi*. It presents a zinc finger CCCH motif and a proline rich motif. The tcZFP2 proteins interact with tcZFP1 via the WW domain and the proline-rich motif, respectively [11]. Since they are expressed in different life-cycle stages, they may allow for a possible modular control of post-transcriptional regulation depending on the parasite stage [12,14]. In fact, It is known that the three *T. brucei* ZFPs were involved in morphological changes and differentiation of the parasite [13,15].

Interestingly, tcZFP1b is expressed only in trypomastigotes (bloodstream stage) suggesting a stage-specific function.

Our results showed that ectopic expression in epimastigotes (insect stage) led to cell replication arrest with incomplete cytokinesis and subsequent cell death. The observed phenotype is almost identical to that obtained when epimastigotes were treated with taxol (a microtubule stabilizing drug). Treatment using as low as 0.1uM taxol inhibited the growth curve in a very similar way as the tetracycline induction of tcZFP1b ectopic expression [18,27]. In accordance, transmission electron micrographs of thin sections of both monster phenotypes looked almost identical. They also showed chromatin in dense granules attached to the nuclear envelope suggesting an arrest in G2 phase of the cell cycle [19,28].

Comparison between the taxol treatment and ectopic expression of tcZFP1b is strongly indicative in favor of a similar mode of action. Thus, we could speculate that ectopic overexpression of tcZFP1b could be stabilizing the sub-pellicular microtubule corset and, in turn, blocking cytokinesis [27,29]. Although it is more difficult to speculate on a direct mode of action of tcZFP1b in this event, we argue in favor of an indirect action through one o more intermediates.

It is well known that cytokinesis depends on the dynamics of this sub-pellicular corset. In fact, the microtubule array is cross-linked together and is present throughout the full cell cycle with new microtubules being added and the array being inherited in a semi-conservative manner by the two daughter cells [27,29]. Thus, it is clear that a stabilization of microtubules would end up blocking the cytokinesis process.

The tcZFP1b is an RBP that could regulate a set of targets that, in turn, could potentially regulate another set of targets. In an attempt to understand the whole picture of its mode of action we decided to look genome-wide instead of looking at the specific targets of its RNA binding motif.

An interesting fact observed in the genome-wide analysis was the 2.4X fold downregulation of α-tubulin. Since new microtubules are constantly needed in order to progress to cytokinesis, this downregulation could be one of the hints pointing to the observed cytokinesis arrest. Non-dividing forms, such as trypomastigotes, would have more stabilized microtubules in the sub-pellicular corset than the fast dividing forms (epimastigotes and amastigotes).

Cause or consequence of epimastigotes stopping cell division several hours upon induction was the fact that PGF2 synthase mRNAs were downregulated by 2.6 and 2.5X fold. It is known that production of PGF2α decreased significantly in non-dividing forms [25].

One of the components of the flagellum, paraflagellar rod component par4, was also downregulated 3X fold. It is not clear the function of this component in the flagellum biology but it is tempting to speculate that its downregulation could be linked to the observed phenotype.

Another interesting observation of the genome-wide analysis is the fact that several genes of related functions were upregulated or downregulated in a concerted form. Most of these genes are located in different chromosomes and, thus, in different post-transcriptional units. However, their post-transcriptional levels appear to be concerted.

Notably, several genes involved in amino acid catabolism that use pyridoxal phosphate as cofactor were upregulated between 13X and 3X fold while the pyridoxal kinase mRNA was concomitantly up by 11X fold.

Several genes related to the glyoxylate and dicarboxylate metabolism were also upregulated. Interestingly, amino acid catabolism is linked to the former metabolism through the production of hydroxypyruvate and glyoxylate.

One important conclusion obtained from this work is that post-transcriptional regulons are evident in *T. cruzi*, as it also seems to emerge from other works in *T. brucei* [30,31]. The finding of common sequence motifs in the 3′UTRs reinforces the idea of the regulons model. In this work, we found two different high-confidence motifs, up-h12 in the upregulated and down-h12 in the downregulated genes. Both motifs presented conserved sequence cores exposed in loops. Interestingly, down-h12 resembles a classical AU-rich (ARE) element previously involved in the instability of mRNAs in *T. cruzi* [9]. The list of down-h12 containing genes included fatty acid elongase, fatty acid desaturase, ATPase beta subunit, cytochrome C oxidase and amastin among others (Table S2)

Important to note is that most of the upregulated genes belonged to the family of trans-sialidase like members, a family expressed almost exclusively in trypomastigotes, the parasite form that interact with the mammalian host. The expression of trans-sialidase pseudogenes was also evident. Expression of pseudogenes was shown to have a role in gene expression via the RNA interference (RNAi) system in *T. brucei* [32]. However, since RNAi is lacking in *T. cruzi* [33,34], the upregulation of these pseudogenes remains puzzling.

A general view of the genome-wide analysis seemed to point to the activation of at least part of a program to differentiate to trypomastigotes, although the sole overexpression of tcZFP1b might not be sufficient to accomplish the task. Upregulation of proline racemase (7.1X fold) is remarkable in this context since it was shown that its ectopic expression in epimastigotes resulted in enhanced differentiation to trypomastigotes and enhanced infectivity [22,23].

In addition, the upregulation of several trans-sialidase like mRNAs, the possible stabilization of the microtubule sub-pellicular corset to enter a non-dividing form, the upregulation of a mechanism of energy production through the glycosome and amino acid catabolism, and the possible downregulation of mitochondrial functions are compelling evidence pointing towards that direction.

With this genome-wide analysis in hand, one of the challenging tasks for the near future would be to dissect the chain of events unleashed after overexpression of tcZFP1b in epimastigotes, beginning by looking for direct mRNA targets of its zinc-finger motif. Since the tcZFP2 proteins interact with tcZFP1b, it will be interesting also to look for direct mRNA targets of their zinc-fingers as well.

## Materials and Methods

### Trypanosome cultures and tetracycline induction of pTcINDEX

For inducible expression of tcZFP1b in the parasite, we first generated a cell line expressing T7 RNA polymerase and tetracycline repressor genes by transfecting epimastigotes with the plasmid pLew13 by electroporation as previously described [35]. Stable transfectants were selected and grown in brain-heart-tryptose (BHT) medium supplemented with 10% inactivated fetal calf serum (FCS) and 200 µg/ml G418 (Gibco). This cell line was then transfected with pTcINDEX construct [17] carrying tcZFP1B gene or with the GFP gene, and transgenic parasites were obtained after selection with 200 µg/ml G418 and 200 µg/ml hygromicin B (Calbiochem). Epimastigote cultures were grown to reach a cell density of 5 × 10^6^ parasites/ml and protein expression was induced by the addition of 5-µg/ml of tetracycline for 60-72 hours. Epimastigotes (1 × 10^9^ cells) were harvested by centrifugation at 1,000 x g for 5 min, washed twice in PBS and lysed on ice by incubation with Laemmli’s sample buffer (for Western Blot) or fixed by 4% paraformaldehyde (for indirect immunofluorescence)

### Microscopy Analysis

Confocal microscopy, GFP detection and indirect immunofluorescence (IFI) analysis were done as previously described [16].

For Transmission Electron Microscopy (TEM), 10^7^ epimastigotes were fixed in 2.5% glutaraldehyde in 0.1 M phosphate buffer, pH 7.2, for 60 minutes, washed in the same buffer, post-fixed in 1% OsO_4_ and 0.8% potassium ferrocyanide in 0.1 M sodium cacodylate buffer at room temperature for 40 minutes, washed in 0.1 M phosphate buffer, dehydrated in acetone, and embedded in Polybed resin. Ultrathin sections were stained with uranyl acetate and lead citrate and observed using a TEM Philips EM 301 at CMA (Centro de Microscopias Avanzadas, University of Buenos Aires).

### Fluorescence Activated Cell Sorting (FACS) analysis

For FACS analysis, epimastigotes expressing inducible tcZFP1b were compared to a non-induced control. Samples were taken at 70 hours after induction when parasites stopped cell division. A total of 10^7^ epimastigotes were washed twice with PBS and fixed in 500 ml of 70% (v/v) ice cold ethanol/PBS overnight at 4°C. The fixed cells were then resuspended in 500 ml of PBS supplemented with 50 μg/ml propidium iodide, 20 μg/ml RNAse A and 2 mM EDTA in PBS before incubation at 37°C for 30 min. FACS analysis was performed with a Becton Dickinson FACSCalibur using FL2-A (detecting fluorescence emission between 543 and 627 nm, propidium iodide), the forward scatter and the side scatter detectors. A total of 10,000-gated events were harvested from each sample. Data were interpreted using the WinMDI 2.9 software (Scripps Research Institute).

### RNA extraction, processing and Pyrosequencing

Total RNA from 1 × 10^9^ induced and non-induced control epimastigotes were extracted from biological duplicates using standard procedures [35].

Total RNA quantity and quality were assessed using the Agilent 2100 Bioanalyzer (Agilent technologies).

Poly-A+ RNA was selected by oligo-dT chromatography in two rounds (Dynabeads mRNA DIRECT kit, Invitrogen), using the total RNA extracted from biological duplicates for each condition. The two rounds purification of polyA+ allowed diminishing the rRNA contamination below 10%. A total amount of 200 ng RNA was quantitated by Ribogreen and quality assessed on an RNA 6000 Pico Chip on the Agilent 2100 Bioanalyzer. RNA was fragmented using a solution of ZnCl_2_ according to 454 cDNA rapid library preparation method manual, generating fragments with a mean size of 500 bp. Finally, the cDNA was synthesized using random hexamers according to manufacturers instructions (Roche).

The cDNA quality was assessed using a high-sensitivity Chip on the Agilent 2100 Bioanalyzer and subjected to 454 sequencing using standard protocols (Roche) at INDEAR sequencing facility (Rosario, Argentina). A half of the PicoTiter Plate (PTP) was divided in quarters, using one quarter for the biological duplicates of the non-induced condition with two MIDs (Multiplex Identifiers) and the other quarter for the duplicates of induced condition with two MIDs. Raw sequencing data produced 233,310 reads and 206,703 reads for each quarter respectively with median read length of 464 and 455 bases respectively.

### Bioinformatics analysis of transcriptome data

Raw data was filtered for artificial duplicate reads and mapped against the *T. cruzi* CL-Brener esmeraldo and non-esmeraldo haplotypes as references genomes. The mapping was done using the 454 GS Reference Mapper software (Roche). The uniquely mapped reads were taken into account for further processing. Mapping statistics could be found in Fig. S2C.

For normalization purposes, the unique read counts were normalized first by gene length. Since RNA fragmentation produced an average of 500 b, we introduced a factor for each gene as a ratio of gene length to fragment size (500 b). If fragment size is greater than gene length then factor is 1. If fragment size is lower than gene length then a correction factor is introduced in the formula below:

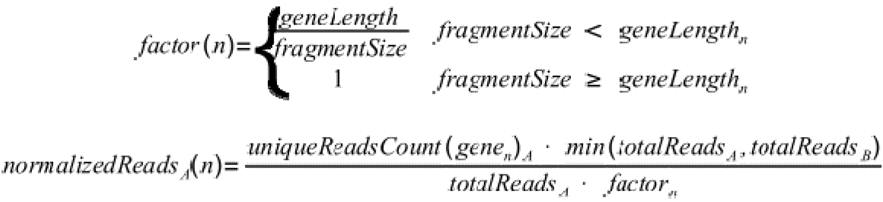

Once normalized reads for conditions A and B were calculated for the biological duplicates, a fold change was calculated as a ratio between induced and non-induced control normalized reads.

The results were parsed into a tabulated spreadsheet format with the GeneDB accession number, fold change, GeneDB description and GO annotation (if available) for each gene (Table S1).

### Computational analysis for sequence motifs search

For 3’ UTR sequence definition, a length of 300 nt downstream to the CDS was used to obtain sequences resembling the 3’UTR, in agreement to previously reported data from trypanosomes [36,37] was downloaded using TcruziDB sequence retrieval tool. Homologue genes within each group with similar 3’UTR were filtered to avoid duplicated sequences. Consensus motifs were predicted from each dataset using CMfinder 0.2 [38]. Candidate motifs obtained were used to build the stochastic context-free grammar (SCFG) model (INFERNAL program). The SCFG for each candidate motif was used to search against the specific data set and the complementary database to obtain the number of hits for each motif (CMSEARCH program). The motif with the highest enrichment in the specific data set was considered to be the best candidate motif. The motif logo was constructed using WebLogo (http://weblogo.berkeley.edu/). Finally, RNAfold server [39] was used to plot the secondary structure of the representative RNA motifs. Differences between groups were examined for statistical significance using chi-square test. Comparison was made between the motif-containing group and random 3’UTR groups (115 lists composed by 50 randomly selected sequences).

## Acknowledgments

The authors wish to thank Dr. Guillermo Alonso, Dr. Claudio Pereira, Dr. Maria Teresa Tellez-Iñon, Dr. Martin Edreira and Dr. Ken Kobayashi for helpful discussions during the writing of this manuscript. The authors thank John Kelly and Martin Taylor (London School of Tropical Medicine, London, United Kingdom) for kindly providing the pTcINDEX expression vector and Dr. Juan José Cazzulo (IIB-INTECH, Buenos Aires, Argentina) for support. JGDG and MPV are members of the career of scientific investigator of CONICET, Argentina.

---

**Figure S1 – FACS analysis and DNA content of normal and monster cells**

**A:** FACS analysis. Upper panel, histogram analysis of control and induced cells. Lower panel, two-dimensional dot plot analysis of the corresponding histogram analysis shown above. The ploidies of the peaks and dots are shown. Epimastigotes were prepared 60hs after induction and fixed. **B:** Confocal microscope analysis of the FACS samples. Left panel shows normal cell division of non-induced sample, right panel shows arrested cytokinesis in induced sample.

**Figure S2 – 454 transcriptome sampling and metrics**

**A:** Epimastigotes growth curve for non-induced (control) and induced cell lines. An arrow indicates the time point of sampling for RNA-seq analysis. **B:** Reference mapping metrics for 454 reads obtained by pyrosequencing for Control and Induced samples. Uniquely mapped 1 and uniquely mapped 2 refers to the biological duplicates reads. Unmapped reads are reported as total for condition. **C:** Overexpression of tcZFP1b after tetracycline induction. The endogenous non-induced tcZFP1a is reported for comparison purposes.

**Figure S3 – 454 Transcriptome sequence coverage and depth.**

**A:** tcZFP1b coverage and SNPs detected. SNPs are indicated above and correspond to differences between *T. cruzi* CL-Brener strain (reference genome) and the *T. cruzi* I strain used in this study; aa, amino acids; nt, nucleotides; syn, synonyms. **B:** High expressed gene coverage example. **C:** Low expressed gene coverage example. Darker reads indicate forward direction. Lighter reads indicate reverse direction.

**Figure S4 – Validation of RNA-seq using qPCR**

Real-time PCR (qPCR) was used to validate RNA-seq results on selected genes. Measurements are the results of triple technical replicates for each biological duplicate. AATc, AATm, Aspartate aminotransferase cytoplasmic and mitochondrial respectively (Tc00.1047053503841.70, Tc00.1047053510945.70); Piryk, pyridoxal kinase (Tc00.1047053507925.40); CPEP, carboxypeptidase (Tc00.1047053504153.160); GAPDH, gliceraldehide-3-P dehydrogenase; AcylCar, acyl carrier protein, mitochondrial precursor (Tc00.1047053511867.140); L35A, ribosomal protein L35A (Tc00.1047053506559.470); PGF2A, prostaglandine F2α synthase (Tc00.1047053507617.9); ATUB; α-tubulin (Tc00.1047053411235.9); ATPase, mitochondrial ATPase beta subunit (Tc00.1047053509233.180).

**Table S1 – Complete 454 transcriptome data for control and induced epimastigotes**

Results obtained from the 454 GS Reference Mapper software and normalized using a custom script as indicated in Materials and Methods were parsed to a tabulated spreadsheet format. The term f1 refers to the Control condition and f2 to the induced condition. Gene Ontology (GO) annotation was included when available.

**Table S2 – Presence of up-h12 and down-h12 motifs in the upregulated and downregulated genes.**

The gene ID, description, motif presence and position in their respective 3’UTRs are indicated. Positions indicate distances from stop codon.

## References

1 Clayton C (2002) Life without transcriptional control? From fly to man and back again. EMBO J 21: 1881–1888.

2 Clayton C, Shapira M (2007) Post-transcriptional regulation of gene expression in trypanosomes and leishmanias. Mol Biochem Parasitol 156: 93–101.

3 Martínez-Calvillo S, Yan S, Nguyen D, Fox M, Stuart K, et al. (2003) Transcription of *Leishmania major* Friedlin chromosome 1 initiates in both directions within a single region. Mol Cell 11: 1291–1299.

4 Worthey E, Martinez-Calvillo S, Schnaufer A, Aggarwal G, Cawthra J, et al. (2003) *Leishmania major* chromosome 3 contains two long convergent polycistronic gene clusters separated by a tRNA gene. Nucleic Acids Res 31: 4201–4210.

5 Liang X, Haritan A, Uliel S, Michaeli S (2003) trans and cis splicing in trypanosomatids: mechanism, factors, and regulation. Eukaryot Cell 2: 830– 840.

6 De Gaudenzi J, Frasch AC, Clayton C (2005) RNA-binding domain proteins in Kinetoplastids: a comparative analysis. Eukaryot Cell 4: 2106–2114.

7 Kramer S, Carrington M (2011) Trans-acting proteins regulating mRNA maturation, stability and translation in trypanosomatids. Trends Parasitol 27: 23–30.

8 Kramer S, Kimblin NC, Carrington M (2010) Genome-wide in silico screen for CCCH-type zinc finger proteins of *Trypanosoma brucei*, *Trypanosoma cruzi* and *Leishmania major*. BMC Genomics 11: 283.

9 Cassola A, De Gaudenzi J, Frasch AC (2007) Recruitment of mRNAs to cytoplasmic ribonucleoprotein granules in trypanosomes. Mol Microbiol 65: 655–670.

10 Caro F, Bercovich N, Atorrasagasti C, Levin MJ, Vázquez MP (2006) *Trypanosoma cruzi*: analysis of the complete PUF RNA-binding protein family. Exp Parasitol 113: 112–124.

11 Caro F, Bercovich N, Atorrasagasti C, Levin MJ, Vazquez MP (2005) Protein interactions within the TcZFP zinc finger family members of *Trypanosoma cruzi*: implications for their functions. Biochem Biophys Res Commun 333: 1017–1025.

12 Hendriks EF, Robinson DR, Hinkins M, Matthews KR (2001) A novel CCCH protein which modulates differentiation of *Trypanosoma brucei* to its procyclic form. EMBO J 20: 6700–6711.

13 Hendriks E, Matthews K (2005) Disruption of the developmental programme of *Trypanosoma brucei* by genetic ablation of TbZFP1, a differentiation-enriched CCCH protein. Mol Microbiol 57: 706–716.

14 Paterou A, Walrad P, Craddy P, Fenn K, Matthews K (2006) Identification and stage-specific association with the translational apparatus of TbZFP3, a CCCH protein that promotes trypanosome life-cycle development. J Biol Chem 281: 39002–39013.

15 Ling AS, Trotter JR, Hendriks EF (2011) A Zinc Finger Protein, TbZC3H20, Stabilizes Two Developmentally Regulated mRNAs in Trypanosomes. J Biol Chem 286: 20152–20162.

16 Westergaard GG, Bercovich N, Reinert MD, Vázquez MP (2010) Analysis of a nuclear localization signal in the p14 splicing factor in *Trypanosoma cruzi*. Int J Parasitol, 40: 1029–1035.

17 Taylor M, Kelly J (2006) pTcINDEX: a stable tetracycline-regulated expression vector for *Trypanosoma cruzi*. BMC Biotechnol 6: 32–32.

18 Baum SG, Wittner M, Nadler JP, Horwitz SB, Dennis JE, et al. (1981) Taxol, a microtubule stabilizing agent, blocks the replication of *Trypanosoma cruzi*. Proc Natl Acad Sci USA 78: 4571–4575.

19 Elias MCQB, Faria M, Mortara RA, Motta MCM, de Souza W, et al. (2002) Chromosome localization changes in the *Trypanosoma cruzi* nucleus. Eukaryot Cell 1: 944–953.

20 Wang Z, Gerstein M, Snyder M. (2009) RNA-Seq: a revolutionary tool for transcriptomics. Nat Rev Genet 10: 57–63.

21 Frasch AC (2000) Functional diversity in the trans-sialidase and mucin families in *Trypanosoma cruzi*. Parasitol today 16: 282–286.

22 Coutinho L, Ferreira MA, Cosson A, Batista MM, Batista DDGJ, et al. (2009) Inhibition of *Trypanosoma cruzi* proline racemase affects host-parasite interactions and the outcome of in vitro infection. Mem. Inst. Oswaldo Cruz 104: 1055–1062.

23 Chamond N, Goytia M, Coatnoan N, Barale JC, Cosson A, et al. (2005) *Trypanosoma cruzi* proline racemases are involved in parasite differentiation and infectivity. Mol Microbiol 58: 46–60.

24 Urbina JA, Osorno CE, Rojas A. (1990) Inhibition of phosphoenolpyruvate carboxykinase from *Trypanosoma* (Schizotrypanum) *cruzi* epimastigotes by 3-mercaptopicolinic acid: in vitro and in vivo studies. Arch. Biochem. Biophys 282: 91–99.

25 Kubata BK, Duszenko M, Kabututu Z, Rawer M, Szallies A, et al. (2000) Identification of a novel prostaglandin F(2alpha) synthase in *Trypanosoma brucei*. J. Exp. Med 192: 1327–1338.

26 Coughlin BC, Teixeira SM, Kirchhoff LV, Donelson JE (2000) Amastin mRNA abundance in *Trypanosoma cruzi* is controlled by a 3’-untranslated region position-dependent cis-element and an untranslated region-binding protein. J Biol Chem 275: 12051–12060.

27 Hammarton T, Monnerat S, Mottram J (2007) Cytokinesis in trypanosomatids. Curr Opin Microbiol 10: 520–527.

28 Elias M, Dacunha J, Defaria F, Mortara R, Freymuller E, et al. (2007) Morphological Events during the *Trypanosoma cruzi* Cell Cycle. Protist 158: 147–157.

29 de Souza W (2009) Structural organization of *Trypanosoma cruzi*. Mem. Inst. Oswaldo Cruz 104 Suppl 1: 89–100.

30 Archer SK, Inchaustegui D, Queiroz R, Clayton C (2011) The Cell Cycle Regulated Transcriptome of *Trypanosoma brucei*. PLoS ONE 6: e18425

31 Archer SK, Luu VD, De Queiroz R., Brems S, Clayton C, et al. (2009) *Trypanosoma brucei* PUF9 Regulates mRNAs for Proteins Involved in Replicative Processes over the Cell Cycle. PLoS Pathogens 5: e1000565

32 Wen YZ, Zheng LL, Liao JY, Wang MH, Wei Y, et al. (2011) Pseudogene-derived small interference RNAs regulate gene expression in African *Trypanosoma brucei*. Proc Natl Acad Sci USA 108: 8345–8350.

33 Ullu E, Tschudi C, Chakraborty T (2004) RNA interference in protozoan parasites. Cell. Microbiol 6: 509–519.

34 DaRocha WD, Otsu K, Teixeira SMR, Donelson JE (2004) Tests of cytoplasmic RNA interference (RNAi) and construction of a tetracycline-inducible T7 promoter system in *Trypanosoma cruzi*. Mol Biochem Parasitol,133: 175–186.

35 Ben-Dov CP, Levin MJ, Vázquez MP (2005) Analysis of the highly efficient pre-mRNA processing region HX1 of *Trypanosoma cruzi*. Mol Biochem Parasitol 140: 97–105.

36 Brandão A, Jiang T (2009). The composition of untranslated regions in *Trypanosoma cruzi* genes. Parasitol Int 58: 215–219.

37 Benz C, Nilsson D, Andersson B, Clayton C, Guilbride DL (2005). Messenger RNA processing sites in *Trypanosoma brucei*. Mol Biochem Parasitol 143: 125–134.

38 Yao Z, Weinberg Z, Ruzzo WL (2006) CMfinder—a covariance model based RNA motif finding algorithm. Bioinformatics 22: 445–452.

39 Hofacker IL (2003) Vienna RNA secondary structure server. Nucleic Acids Res 31: 3429–3431.

